# ATP:Mg^2+^ shapes condensate properties of rRNA-NPM1 *in vitro* nucleolus model and its partitioning of ribosomes

**DOI:** 10.1101/2021.12.22.473778

**Authors:** N. Amy Yewdall, Alain A. M. André, Merlijn H. I. van Haren, Frank H.T. Nelissen, Aafke Jonker, Evan Spruijt

**Affiliations:** Institute for Molecules and Materials, Radboud University, Heyendaalseweg 135, 6525 AJ, Nijmegen, the Netherlands

**Keywords:** Condensate, Nucleolus, Gelation, Magnesium, ATP, RNA

## Abstract

Nucleoli have viscoelastic gel-like condensate dynamics that are not well represented *in vitro*. Nucleoli models, such as those formed by nucleophosmin 1 (NPM1) and ribosomal RNA (rRNA), exhibit condensate dynamics orders of magnitude faster than *in vivo* nucleoli. Here we show that an interplay between magnesium ions (Mg^2+^) and ATP governs rRNA dynamics, and this ultimately shapes the physical state of these condensates. Using quantitative fluorescence microscopy, we demonstrate that increased RNA compaction occurs in the condensates at high Mg^2+^ concentrations, contributing to the slowed RNA dynamics. At Mg^2+^ concentrations above 7 mM, rRNA is fully arrested and the condensates are gels. Below the critical gel point, NPM1-rRNA droplets age in a temperature-dependent manner, suggesting that condensates are viscoelastic materials, undergoing maturation driven by weak multivalent interactions. ATP addition reverses the dynamic arrest of rRNA, resulting in liquefaction of these gel-like structures. Surprisingly, ATP and Mg^2+^ both act to increase partitioning of NPM1-proteins as well as rRNA, which influences the partitioning of small client molecules. By contrast, larger ribosomes form a halo around NPM1-rRNA coacervates when Mg^2+^ concentrations are higher than ATP concentrations. Within cells, ATP levels fluctuate due to biomolecular reactions, and we demonstrate that a dissipative enzymatic reaction can control the biophysical properties of *in vitro* condensates through depletion of ATP. This enzymatic ATP depletion also reverses the formation of the ribosome halos. Our results illustrate how cells, by changing local ATP concentrations, may regulate the state and client partitioning of RNA-containing condensates such as the nucleolus.

**Significance Statement:**

- There is a significant discrepancy between the dynamics of *in vitro* nucleolus models and *in vivo* nucleoli – with the latter more gel-like.
- The interplay between Mg^2+^ ions, ATP and the nucleolus components – specifically RNA – governs the dynamics, and ultimately the physical state, of nucleolus-like condensates.
- We show that the nucleolus are dynamically adapting condensates, responding to local ATP concentrations through Mg^2+^-induced compaction of the RNA, and reversible relaxation when ATP binds Mg^2+^ again. Other condensates containing RNA probably respond in similar ways to Mg^2+^ and ATP.

## Introduction

The nucleolus is a liquid-liquid phase-separated subcompartment formed during eukaryotic interphase as the site of ribosome biogenesis. Consequently, ribosomal RNA (rRNA) and various RNA-binding proteins are predominantly found in this dynamic, but highly viscous, condensate. The gel-like dynamics of *in vivo* nucleoli, with slow fusion time scales of ∼30 mins and non-spherical shapes (1-3), is not well represented by *in vitro* models, which are often liquid-like. The granular component of the nucleolus can be reconstituted *in vitro* using nucleophosmin 1 (NPM1) protein and rRNA (2). The contribution of NPM1 proteins to condensate properties is well characterized, with fast FRAP recovery times ∼7-20 seconds (4, 5), but this is still orders of magnitude separated from *in vivo* nucleoli dynamics. Therefore, we want to consider the RNA component as this was also shown to play an important role in shaping the dynamics of other condensates (6, 7).

At the core of nucleoli, rRNA is transcribed in the border region between the fibrillar center and the dense fibrillar component, and a recent elegant study from Brangwynne and co-workers has shown that this pre-rRNA traverses the nucleoli radially outward as it matures (8). The outer region of the nucleoli, in the so-called granular component, the rRNA is folded around the pre-ribosomal protein subunits with the aid of various protein chaperones (9, 10). Once folded, the pre-ribosomes are thermodynamically ejected out of the nucleolus (11). rRNA is a fundamental component of nucleoli (9, 10). rRNAs are large, complex polymers that fold into secondary structures at precise magnesium ion (Mg^2+^) concentrations (12, 13). The intracellular availability of free Mg^2+^ can range from 0.5-1 mM in eukaryotic cells and 1-5 mM in bacterial cells, with *in vivo* concentrations of Mg^2+^ bound to various metabolites ranging between 20-100 mM (13, 14). Additionally, the free Mg^2+^ concentrations can vary throughout the cell cycle due to changes in levels of adenosine triphosphate (ATP), a nucleotide that strongly complexes Mg^2+^ (15-17).

Despite this, there has yet been no systematic analysis of the effect of Mg^2+^ and ATP on RNA compaction and this effect on condensate properties. In this work, we examined the interplay of Mg^2+^ and ATP on an *in vitro* reconstituted nucleolus, in order to shed light on how these small molecules can reversibly affect RNA compaction to shape the properties of nucleolus-like condensates, as well as to influence client molecule partitioning. In particular, we explored the intriguing formation of a 70S ribosome halo that transiently associated with the NPM1-rRNA condensates, reminiscent of the *in vivo* function of these condensates. Together, our results not only help explain the discrepancies between *in vitro* and *in vivo* nucleoli dynamics, but also show that nucleoli are adaptive condensates, responding to local ATP concentrations through reversible Mg^2+^-induced compaction of the RNA component. Transient changes to ATP levels may be how cells regulate the amount of free Mg^2+^ and therefore how they can control the dynamics of RNA-based condensates, such as the nucleolus.

## Results

### Mg^2+^-induced RNA compaction slows dynamics and leads to gelation

In order to obtain a better understanding of why *in vivo* nucleoli are so slow at fusing, we used NPM1-rRNA model condensates to first assess the effect of Mg^2+^ ions on the dynamics of each component (**Figure 1A**). Simultaneous fluorescence after photobleaching (FRAP) analyses for both protein and RNA components demonstrated a noticeable difference in FRAP recovery even at 0 mM Mg^2+^ concentration, with a faster recovery half-life (τ) for NPM1 at 18.3 ± 2.4 s and slower recovery for rRNA at 30.7 ± 6.1 s (**Figure 1B and C**). The τ for NPM1 is comparable to *in vivo* values of around 20 s (2, 4). The difference between protein and RNA recoveries is reminiscent of other intracellular protein-RNA condensates, where the constituent RNAs recover slower than protein components; this was attributed to multivalent RNA-RNA interactions (6, 7). In order to promote such RNA-RNA interactions in the NPM1-rRNA condensates, we increased the Mg^2+^ concentrations and observed the NPM1 protein and RNA dynamics diverge, with rRNA recovering slower and to a lesser extent (**Figure 1D and E**). At Mg^2+^ concentrations higher than 7 mM, rRNA is fully arrested with τ >1500 s and percentage recoveries decreased to <10% (**Figure 1E**). These findings suggest a gradual transition from liquid to an arrested, gel state of the RNA component of the nucleolus, which is supported by the scaling behavior of the relaxation times near the critical point (**Figure 1D**) (18, 19). The NPM1 component remained astonishingly mobile within the rRNA gel network as the Mg^2+^ concentration increased, with low τ (**Figure 1D**), which is consistent with other reported FRAP recoveries (2, 4). The percentage recovery of NPM1 proteins appeared to not significantly decrease until 20 mM Mg^2+^ was reached (**Figure 1E**). NPM1 thus acts as a shuttle between different regions of the gelled RNA network. Finally, we note that when rRNA dynamics is fully arrested, the condensates adopt a fractal morphology (**Figure 1F**) with gel-like states that can still slowly fuse together within the minute’s time-scale (**Video S1**), reminiscent of nucleoli fusing *in vivo*. These results suggest that RNA-RNA interactions play an important role in shaping condensate dynamics.

**Figure 1.**
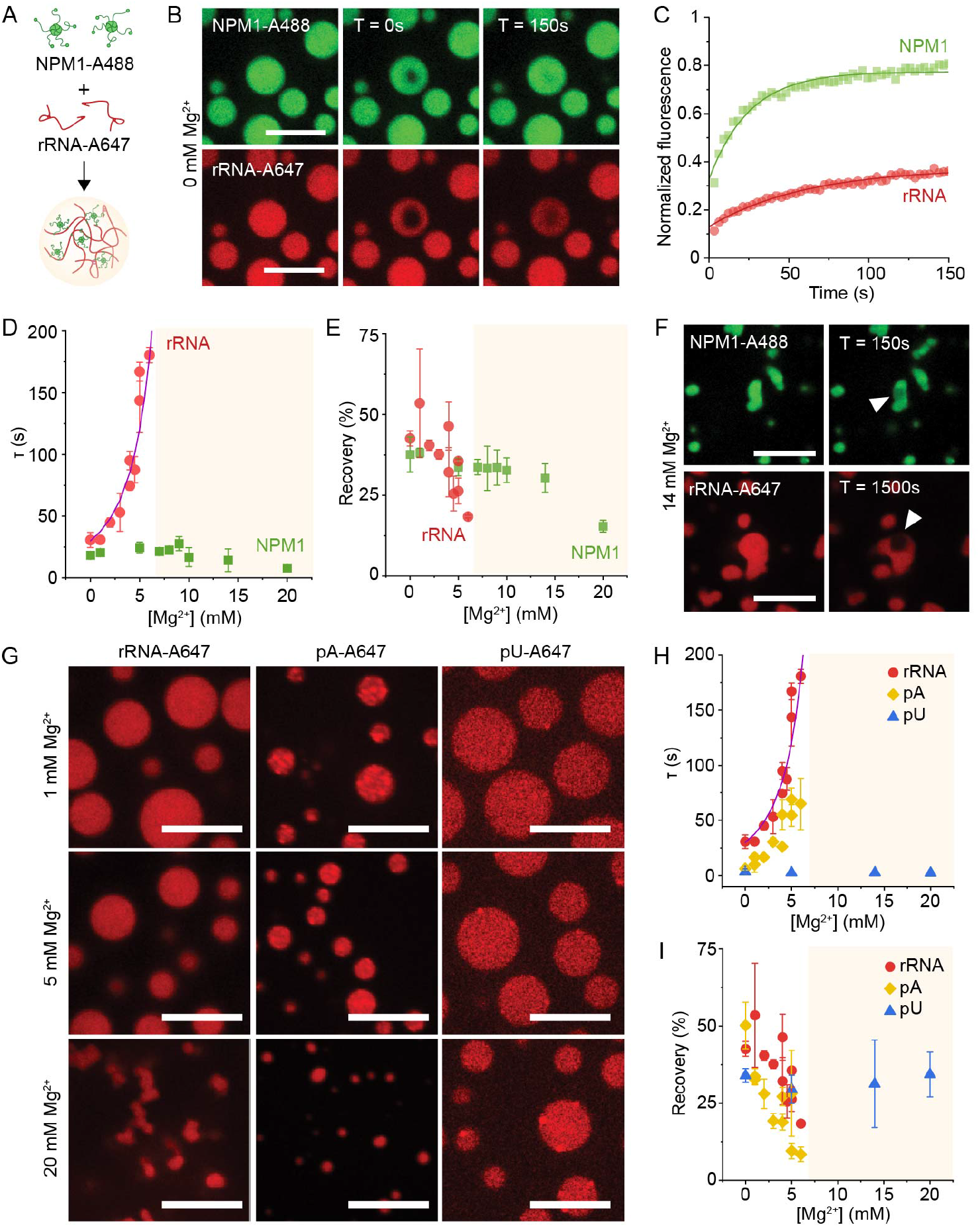
Mg^2+^-induced RNA compaction leads to slow dynamics and gelation. (**A**) NPM1 protein and rRNA, both labelled with different fluorophores, are mixed together to form condensates that show differences in FRAP recovery (**B & C**), where NPM1 (green) recovers faster than rRNA (red). (**D**) The recovery of rRNA slows to a halt at Mg^2+^ ion concentrations > 7 mM (shaded in yellow) and shows critical scaling behavior (purple fitted line) indicating it forms a gel, whereas the NPM1 protein remains mobile. (**E**) The decrease in percentage recovery for both NPM1 and rRNA reflects the gel environment at higher Mg^2+^ concentrations. (**F**) These droplets become fractal gels at higher Mg^2+^ where the rRNA is fully arrested, indicated by bleached regions not recovering (white arrows). In order to test the RNA compaction hypothesis, NPM1-rRNA condensate morphology (**G**) and FRAP recovery parameters (**H & I**) were compared with NPM1-pA and NPM1-pU condensates. The errors in this figure are standard deviations from triplicate measurements. Scale bars are all 10 μm.

Mg^2+^ stabilization of RNA-RNA interactions and corresponding compaction of RNA chains is well reported (13, 17). The observed rRNA arrest at high Mg^2+^ is likely due to the compaction of RNA, promoted by enhanced RNA-RNA base-paring and stacking interactions. To further explore this process, NPM1 condensates were made with other homopolymeric RNAs with different propensities for RNA-RNA interactions. Poly-adenosine (pA) RNA is known to form base-stacking interactions with Mg^2+^, whereas poly-uridine (pU) remains largely unstructured (20). At 20 mM Mg^2+^, condensates made from rRNA or pA both formed fractal gels, whereas condensates made from pU remained round, liquid droplets (**Figure 1G**) that still readily fused with one another within seconds (**Video S2**). The FRAP parameters also reflected this, with τ increasing for pA at increasing Mg^2+^, but pU τ’s were unaffected by Mg^2+^ (**Figure 1H**).

The gelation induced by RNA compaction can also be seen in decreasing percentage recoveries for pA at increasing Mg^2+^ concentrations, while pU percentages remained relatively unchanged (**Figure 1I)**, in agreement with previous studies on Mg^2+^-induced folding of single 16S rRNA and pU mRNA molecules (21). Together, these results suggest that RNA compaction is facilitated by Mg^2+^ induced RNA-RNA interactions, which occur in RNAs with strong base pairing and base stacking interactions, such as rRNA and pA, but not in RNAs with weak base stacking interactions, such as pU.

### Tuning rRNA compaction with temperature and ATP

We hypothesize that rRNA forms a viscoelastic network in the *in vitro* nucleoli, based indirectly on the partial FRAP recoveries and slowing dynamics with increasing Mg^2+^. Additional evidence for the viscoelastic nature of the rRNA component is its gradual aging to a more structured state, with slower relaxation observed as an increasing τ over time that reaches a plateau (**Figure 2A**). In contrast, the τ of NPM1 remains relatively constant over time (**Figures 2B and S1**). We also showed that this relaxation varies non-linearly with temperature (T), which also supports the viscoelastic nature of rRNA in the condensates. In a pure viscous liquid, where FRAP recovery is commonly attributed to simple Stokes-Einstein diffusion, the diffusion coefficient should increase as 1/T, and the recovery time should be linearly proportional to temperature. However, this is clearly not the case as there is a strong non-linear dependence of τ on T (**Figure 2B**). A closer inspection reveals that the temperature dependence shows signs of an activated process that leads to structuring and relaxation. In fact, our data can be fitted using an Arrhenius equation with a fixed activation energy for temperatures above the gel point temperature (**Figure S2A**). Below that point, the activation energy increases to a higher level, as shown by the transistivity plot (**Figure S2B**), which is characteristic of gels and glasses (22). Therefore, even at 0 mM Mg^2+^, the rRNA network appears to be a viscoelastic material, held together by multivalent intra- and intermolecular RNA-RNA interactions. In contrast, the NPM1 protein can diffuse freely as a chaperone under all conditions. This is consistent with observations that nucleoplasmic NPM1 concentrations increase with higher overall NPM1 expression levels (11).

**Figure 2.**
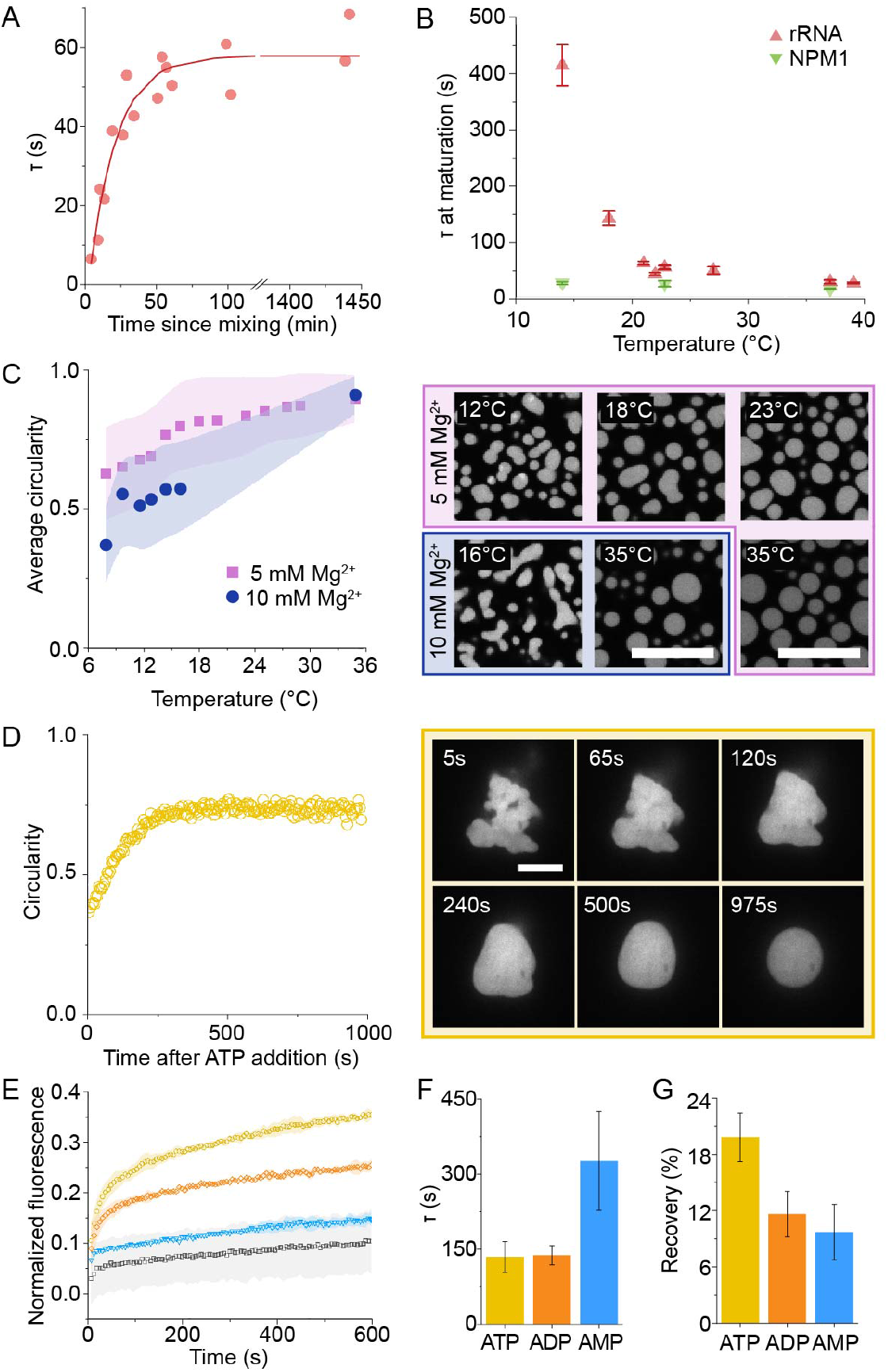
Temperature and ATP can reverse the effect of Mg^2+^-induced RNA condensation. (**A**) NPM1-rRNA condensates aging at 20°C with individual τ for rRNA measured over time (dots) and a fitted exponential curve (line). (**B**) The extracted plateau τ at maturation for rRNA (red) and NPM1 (green) was plotted for aged droplets at different temperatures in 0 mM Mg^2+^ buffer. The errors here are derived from the exponential fits. (**C**) Forming NPM1-rRNA condensates at 8°C in buffer containing magnesium resulted in irregular gel-like morphologies that changed to round droplets at increasing temperatures, with increased circularities. (**D**) The fractal gels formed in 14 mM Mg^2+^ buffer also liquefied after 11 mM ATP addition, with increasing circularity reflected in the changed morphology of the NPM1-rRNA condensates. (**E**) The FRAP recovery over times for samples that contain ATP (yellow), ADP (orange) or AMP (blue) compared with just the 5 mM Mg^2+^ buffer (grey). The extracted τ (**F**) and percentage recoveries (**G**) indicate that the better Mg^2+^-chelating ability of ATP compared to other adenosine nucleotides caused the condensates to liquefy. The errors in figures E-G are standard deviations from at least duplicate measurements. Scale bars are all 10 μm.

Since Mg^2+^ stabilizes RNA-RNA interactions, it was anticipated that this could affect the τ at maturation of condensates formed in Mg^2+^ buffer. Indeed, condensates made in 5 mM Mg^2+^ had slower observed τ at maturation, which decreased at higher temperature (**Figure S3**). This result made us curious about the effect of temperature on Mg^2+^-stabilized RNA-RNA interactions, and how this could impact condensate morphology. NPM1-rRNA condensates formed at 8°C, in either 5 mM or 10 mM Mg^2+^ buffer after 60 minutes of incubation, had a striking non-spherical morphology (**Figures 2C and S4**). The 10 mM Mg^2+^ condensates appeared smaller and had lower average circularity than the 5 mM Mg^2+^ condensates, but both circularities were lower than at higher temperatures, where values increased closer to 1, which is a characteristic of round liquid droplets(23). These results suggest that RNA is fully arrested at lower temperatures in 10 mM Mg^2+^, with the small gel-like condensates unable to completely fuse together. Instead, the small gel droplets partially fuse to form a fractal gel network similar to other gel-like RNA condensates (6). As temperature was increased, the gel-like condensates relaxed and coalesced, eventually, into round droplets, which resulted in increased average circularity of the condensate population (**Figure 2C**). In fact, the condensates appeared round at 35 °C, with circularities close to 1, for both Mg^2+^ concentrations. This temperature-dependent relaxation of the NPM1-rRNA condensates manifesting as a morphological change provides further evidence for gelled states being caused by RNA compaction that results in incomplete fusion of gel condensates. The adaptive morphology changes of NPM1-condensates within the temperature range of living systems, hints that this could be an important parameter to consider for nucleoli dynamics.

In homeostatic eukaryotic cells, Mg^2+^ concentrations and temperature are often well regulated (24, 25), but local ATP levels are known to fluctuate (15, 16). ATP strongly chelates Mg^2+^, changing the effective free Mg^2+^ concentrations. When we tested the effect of ATP addition to the NPM1-rRNA condensates that were incubated with Mg^2+^, we observed that ATP caused the gel-like morphologies to liquefy, and the circularity increased over 20 minutes (**Figure 2D**). ATP relaxes the rRNA network, with FRAP recoveries corresponding to the remaining free Mg^2+^ concentrations (**Figure S5**). In order to probe the influence of ATP on RNA compaction, we used SYBR Gold fluorescence to probe the RNA-RNA interactions within the condensates. SYBR Gold is a fluorescent dye that intercalates double stranded nucleic acids (26), and we hypothesize that Mg^2+^ will stabilize such interactions between the rRNAs. This was indeed the case, as the average SYBR Gold fluorescence detected inside the NPM1-rRNA condensates increased by ∼41% on Mg^2+^ addition (**Figure S6**). Subsequently, the SYBR Gold fluorescence decreased when ATP was added, providing indirect evidence for decreased RNA compaction due to ATP chelating Mg^2+^ (**Figure S6**). This liquefying effect is less pronounced for other adenosine nucleotides (ADP and AMP) that do not chelate Mg^2+^ as effectively (**Figure 2E-G**). These results demonstrate how ATP can be a key regulator of free Mg^2+^ concentrations that can impact NPM1-rRNA condensate dynamics.

### Consequences of ATP:Mg^2+^ on condensate function: partitioning of clients

Alongside chelating Mg^2+^, ATP can also bind to the nucleic acid interacting domain of NPM1 (27, 28), which could lead to an altered chemical microenvironment inside the condensates. Inspired by the profound changes to condensate dynamics observed with Mg^2+^ and ATP, we were curious whether these small molecules would influence the partitioning coefficients (*Kp*) of component proteins and RNAs inside the condensates. Indeed, we observed that, at higher Mg^2+^, the rRNA *Kp* increased on compaction, and the NPM1 protein *Kp* decreased, as more RNA-RNA interactions could exclude protein-RNA interactions (**Figure 3A and B**). Additionally, ATP increased the *Kp* of both RNA and protein components. We speculate that ATP, when bound to the nucleic acid interacting domain, can also stabilize the interaction between NPM1-rRNA. As a result, a synergistic increase in partitioning of both components inside the condensates occurred when Mg^2+^ and ATP were present.

**Figure 3:**
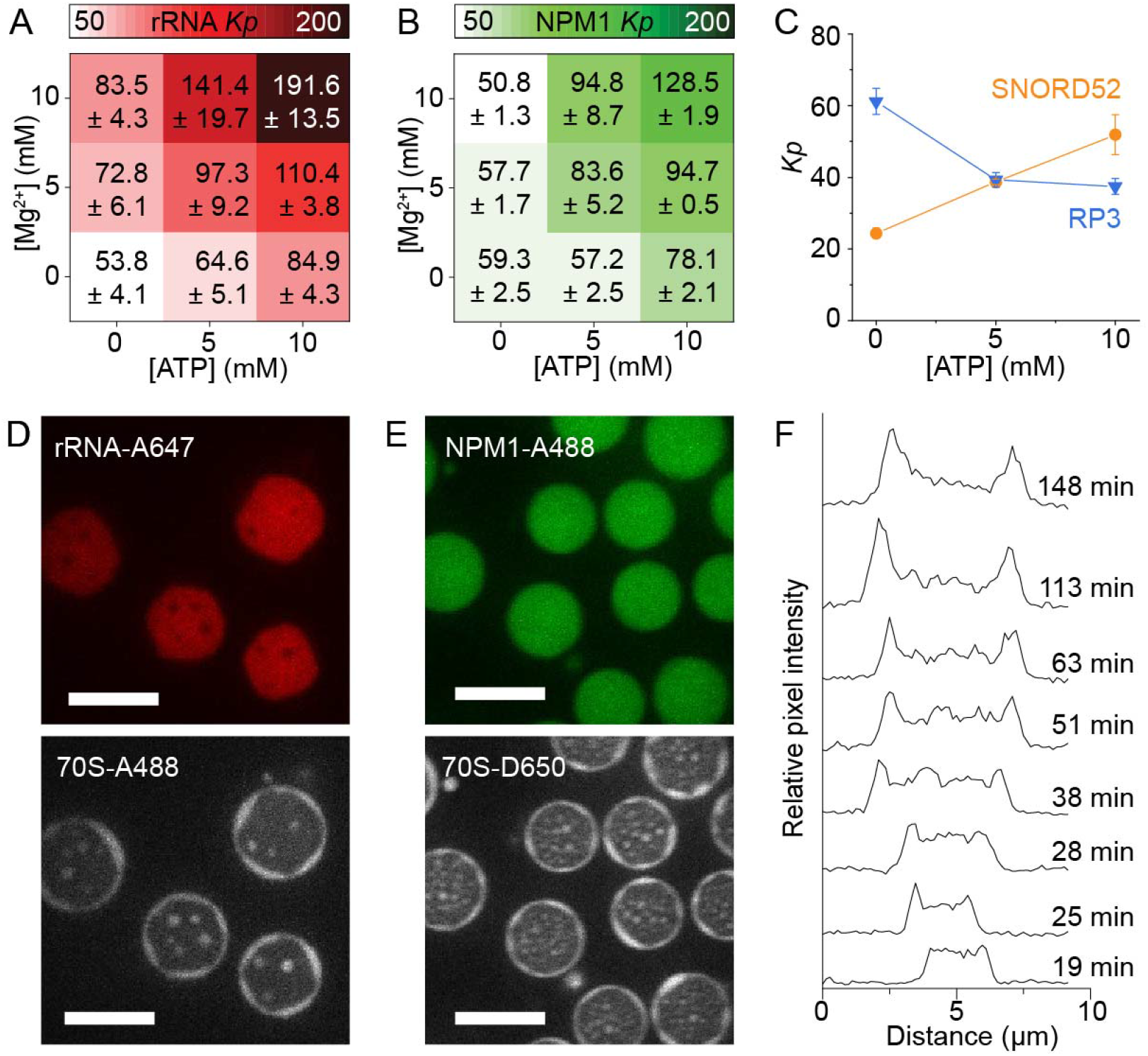
The effect of Mg^2+^ and ATP on partitioning of components, clients and 70S ribosomes. The partitioning coefficients of rRNA (**A**) and NPM1 (**B**) at different Mg^2+^ and ATP concentrations. (**C**) The partitioning coefficients of 5,6-FAM-RP3 peptide (blue) and 5,6-FAM-SNORD52 RNA (orange) at different ATP concentrations in a 5 mM Mg^2+^ base buffer. Confocal microscope images of NPM1-rRNA condensates with 70S ribosome as the client (**D & E**). (**D**) The images show that rRNA-A647 is excluded from the ribosome fluorescence, whereas (**E**) shows NPM1-A488 fluorescence as round droplets that occupy both locations where there is 70S ribosome and rRNA. (**F**) The formation of the ribosome halo over time as indicated by an increase in fluorescence pixel intensity as peaks at the edges of the droplet. The errors in this figure are standard deviations from triplicate measurements. Scale bars are all 10 μm.

We then tested whether the altered microenvironment has a functional consequence in the partitioning of two model clients, the (RRASL)3 peptide (RP3) and the SNORD52 RNA, which interact non-specifically and specifically with NPM1-rRNA condensates, respectively. RP3 resembles arginine rich peptides that can electrostatically interact with both the NPM1 protein and rRNA (29), whereas SNORD52 is a small nucleolar RNA that binds to NPM1 (30) (**Figure 3C**). The *Kp* of both clients followed trends that are expected based on their interaction strengths with the condensate components (**Figure 3C**). The SNORD52 RNA *Kp* increased at higher ATP concentrations as more NPM1 partitioned into the condensates. Conversely, the RP3 *Kp* decreased as less binding surface was expected due to the synergistic partitioning and interaction of NPM1 and rRNA at higher ATP concentrations. These results indicate that Mg^2+^:ATP can be used to tune client partitioning within NPM1-rRNA condensates.

As nucleoli are the sites of ribosome biogenesis, we next investigated the effect of Mg^2+^ and ATP on the partitioning of ribosomes into NPM1-rRNA condensates. Here, we used labelled mature 70S ribosomes that should remain intact at Mg^2+^ concentrations above 5 mM (14). Therefore, it is unsurprising that at increasing Mg^2+^, the *Kp* decreased as the ribosomes remained intact in solution (**Figure S7**). In contrast, ATP addition resulted in a 30% increase in *Kp*, likely due to chelation of Mg^2+^ that was previously bound to the ribosomes, thereby destabilizing the structure and causing it to partition inside the NPM1-rRNA condensate (**Figure S7**). Interestingly, under conditions where Mg^2+^ concentrations exceeded ATP concentrations, the ribosomes formed a striking halo around the NPM1-rRNA condensates (**Figures 3D and E**). The ribosome halo excluded the rRNA, and resulted in a deformed rRNA condensate shape (**Figure 3D**). In contrast, the NPM1 fluorescence is homogeneously distributed throughout the condensates localizing to both the ribosome and rRNA (**Figure 3E**), which is expected from a protein that can bind to the rRNAs present. Indeed, these heterotypic multicomponent interactions of NPM1 with rRNA and the pre-ribosomal subunits, have been suggested to drive ribosome assembly (11), and here we show that NPM1 bridges the interactions between the rRNA component within the condensates and the 70S ribosomes in the halo.

In order to examine our hypothesis that the halo appeared due to partly destabilized 70S ribosomes, we monitored the condensate growth over time. The ribosome halo appeared after ∼25 minutes, with an increased fluorescence intensity observed around the edge of the condensates (**Figure 3F**). Here, the ribosome-NPM1-rRNA condensates exhibited decelerated fusion between larger droplets, in the order of tens of minutes (**Video S3**). A possible reason for slow droplet fusion could be due to the ribosome halo rearrangements that would be necessary for rRNA-NPM1 condensates to fuse. Indeed, we found that the ribosome halo had slow FRAP dynamics with an average τ of 490 ± 10 s, and percentage recoveries of 45 ± 11%. As the droplets fused, the ribosome halo seemed to become incorporated into the condensate and appeared inside as bright spots, with some droplets stalled mid-fusion (**Video S3**). As a control, we made NPM1-rRNA condensates in 10 mM Mg^2+^ and left these to mature for 2 hours, before 5 mM ATP was added, to verify that the observed halo was not a transient phenomenon linked to nucleoli maturation, but rather a result of destabilized ribosomes that originate from the dilute phase. Here, the ribosome halo also appeared after approximately ∼10 minutes of ATP addition, alongside separate droplets composed of NPM1 and ribosomes (**Video S4, Figure S8**), which is reminiscent of *in vivo* experiments showing NPM1 interacting with 60S pre-ribosomes (31). Our results suggest that the ATP addition destabilizes the ribosomes in solution, which in turn could liberate rRNA from the ribosomes to interact with NPM1, forming the halo. The exclusion of partially destabilized ribosomes from the NPM1-rRNA condensates is an exciting result as this corroborates the *in vivo* behavior of nucleoli (11), but to our knowledge this is the first time this phenomenon has been observed *in vitro*. Therefore this work not only provides a useful platform for further studying thermodynamic exclusion from condensates, but also suggests that ATP:Mg^2+^ could be one of the ways cells can alter the condensate environment to drive ribosome formation.

### Enzymatic depletion of ATP levels changes NPM1-rRNA condensate properties and function

Within cells, ATP levels fluctuate due to a variety of enzymes: from ATP-dependent chaperones to reactions that consume ATP (15-17). Here we demonstrate that ATP removal using a dissipative enzymatic reaction can also control the condensate properties of the nucleolus-like condensates made from NPM1 and rRNA. Using apyrase, an enzyme that converts ATP to AMP (**Figure 4A**), we showed that the rRNA recovery of samples without and with apyrase reflected those for the ATP and AMP nucleotides, respectively, similar to observations in Figure 2 (**Figure 4B-D**). Apyrase can effectively liberate Mg^2+^ from ATP, thereby causing rRNA arrest and morphology changes for samples made in high Mg^2+^ concentrations (**Figure 4E-G**). This change in morphology from round liquid-like to irregular-shaped gel-like states is remarkably similar to observations for purified and *in vivo* nucleoli when ATP was depleted (1, 3). For the purified and *in vivo* nucleoli, ATP-dependent chaperones were hypothesized to facilitate liquid-like condensate dynamics. However, our results highlight the impact of liberated Mg^2+^ due to ATP depletion, and demonstrate that changes to free Mg^2+^ levels alone may corroborate these *in vivo* and our *in vitro* observations. Here, enzymes that consume ATP, and not necessarily active ATP-dependent chaperones, are a possible route that cells use to regulate Mg^2+^ levels and impact nucleoli dynamics.

**Figure 4:**
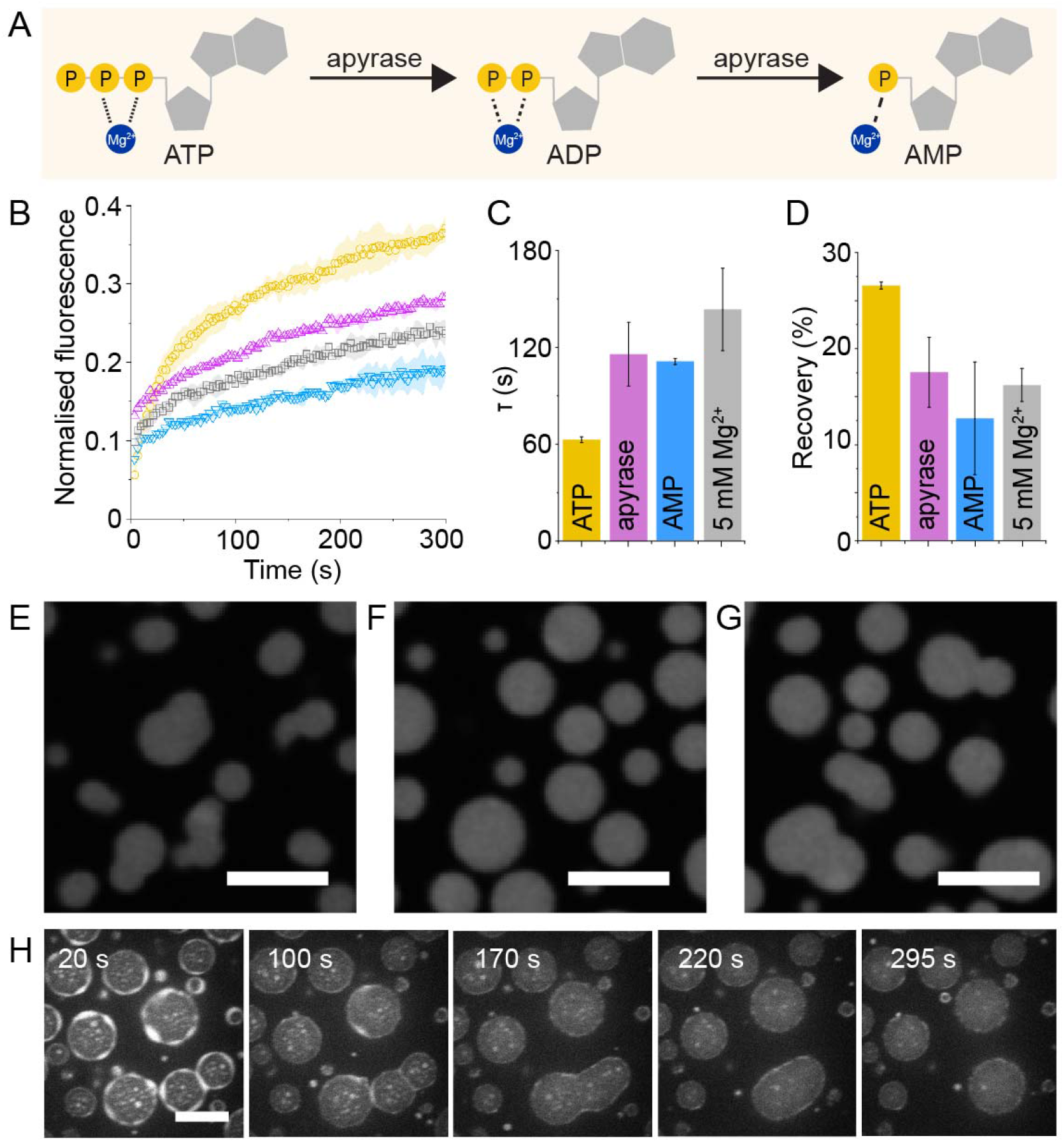
Enzymatic control of ATP concentrations influences condensate morphology and dynamics. (**A**) Apyrase enzymes catalyze the removal of phosphate groups from ATP to form AMP, a nucleotide that poorly chelates Mg^2+^. (**B**) The effect of apyrase on the dynamics of rRNA after FRAP is clearly shown (**B**-**D**) where rRNA recovery in the presence of ATP (yellow) drops when apyrase (purple) is added. In fact, the resulting FRAP parameters (**C & D**) show that apyrase converts ATP to AMP which imbues the rRNA with gel-like dynamics. The errors in this figure are standard deviations from triplicate measurements. The morphology of NPM1-rRNA condensates at 18°C in 5 mM Mg^2+^ buffer (**E**) with 5 mM ATP added (**F**), and when apyrase is also added (**G**) shows the changes in morphology that is expected from gel-like condensates in buffer containing Mg^2+^ that stabilizes RNA-RNA interactions, and round morphology in conditions where ATP chelates the Mg^2+^ and liquefies these interactions. (**H**) This series of confocal images over time of labelled 70S ribosomes show the disappearance of the ribosome halo and puncta, as apyrase was added. ATP depletion results in higher Mg^2+^, which stabilizes the ribosomes and causes them to dissipate back into the dilute phase. Scale bars are all 10 μm.

As nucleoli should exclude fully folded ribosomes (11), we were curious whether ATP removal would affect the previously observed ribosome halo around the NPM1-rRNA condensates. We hypothesized that the ribosome halo was formed from destabilized ribosomes interacting with NPM1, but to what extent the ribosomes were destabilized and whether this was reversible was not yet clear. ATP was depleted using apyrase, which resulted in increased available Mg^2+^ concentrations, and as a result, we observed the striking disappearance of the ribosome halo over time (**Figure 4H**). Additionally, the ribosome-NPM1 puncta formed outside the NPM1-rRNA condensates (**Video S5**) also vanished on apyrase addition, suggesting that fully folded ribosomes are excluded from the NPM1-rRNA condensates. With the disappearance of the ribosome halo, the NPM1-rRNA condensates that were previously stabilized mid-fusion were able to relax and fuse again, and the resulting condensates appeared unexpectedly round in shape (**Video S6**), despite the higher available Mg^2+^ concentrations. The rounder final condensates are hypothesized to be a result of altered Mg^2+^ ions concentrations: although liberated from ATP, the Mg^2+^ will likely bind to the 70S ribosomes making the ion less available to stabilize rRNA-rRNA interactions. These results demonstrate that enzymatic changes to ATP concentrations can have a profound effect on client localization around the NPM1-rRNA condensates, and could indeed be a way how cells change the flux of client molecules interacting with nucleoli.

## Discussion

This work uses *in vitro* reconstituted nucleoli to demonstrate the considerable influence of Mg^2+^:ATP on the dynamics of RNA-based condensates. The irregular morphologies and slow fusion dynamics of nucleoli can be accounted for by changing the ratios of these small molecules alone. For example, the fusion dynamics of NPM1-rRNA condensates at 5 mM Mg^2+^ (**Video S1**) in the order of minutes is reminiscent of *Xenopus* nucleoli (1). The fractal gel morphology, due to incomplete fusion of condensates at high Mg^2+^ is also similar to irregular shaped HeLa nucleoli (3). In additional support of our hypothesis, the ATP depletion studies on purified and *in vivo* nucleoli also resulted in irregularly shaped structures, which was attributed to an unknown ATP-dependent process/enzyme required to maintain nucleoli fluidity (1, 3, 32, 33). However, our ATP-depleted *in vitro* model, when apyrase was added, challenges this perspective. Here, the depleted ATP liberates Mg^2+^, resulting in the morphed gel-like appearance of the nucleoli-like condensates. Our results can help explain the *in vivo* observations without the need for ATP-dependent enzymatic activity to maintain nucleoli fluidity. Of course, nucleoli are much more complex than our model, so at the very least, our data highlight the importance of considering free Mg^2+^ in the interpretation of ATP depletion results. Beyond enzymes, it is also valuable to consider the changes in ATP levels during a cell cycle (16, 34), and how this could impact the fluidity and condensate properties of nucleoli.

RNA compaction is our hypothesis for the underlying mechanism that drives this slowed rRNA dynamics and gel-like morphologies at increased Mg^2+^. We used control experiments with homopolymeric RNAs, as well as SYBR Gold fluorescence, to ascertain that increased RNA compaction at higher Mg^2+^ was indeed observed. Although we have not shown this definitely, the RNA compaction observed is likely due to increased RNA-RNA interactions as this is a well-documented phenomenon at higher Mg^2+^ (12, 13). Our results are also reminiscent of the multivalent RNA-RNA interactions found to drive the formation of gel-like condensates within cells (6). Conversely, when proteins were coerced to make stronger intermolecular interactions via light-induced oligomerization, the resulting condensates were also gel-like in nature (35). Together with these observations, there is a strong indication that rRNA is forming a mesh-like network wherein RNA-RNA interactions are stabilized by Mg^2+^. The changes in material properties due to Mg^2+^ could be a way in which cells control the rRNA flux through the nucleoli. Within this rRNA gel network, the NPM1 (and possibly other proteins) can still diffuse freely.

Interestingly, at low Mg^2+^, we observed aging of the rRNA component of our NPM1-rRNA condensates, which implies it behaves as a viscoelastic liquid. The maturation of the condensates resulted in τ reaching a plateau, the value of which was temperature dependent. This decrease in diffusivity of condensate components, which stabilized after 1h at a constant value, was also observed for *in vitro* protein condensates stabilized by Ni^2+^ interactions (36). Additionally, an aging phenomenon was also observed in *in vivo* TIS11B condensates with slowed FRAP recoveries between 5 and 16 hours (6). Although the physiological relevance is unknown, aging could be a universal feature of viscoelastic condensates formed via multivalent interactions.

Temperature studies on RNA-based condensates have implicated this as an important parameter for inducing the formation of condensates (37, 38). However, there have been very few studies on how temperature affects nucleoli (37, 39), and so far none have shown the temperature-induced changes to morphology of RNA-based condensates, as described in our work. In contrast, temperature responsive gels are abundantly documented in materials science (40). Through this avenue, we used FRAP τ at different Mg^2+^ concentrations to explore the critical scaling that occurs to the RNA component of our NPM1-rRNA condensates at high Mg^2+^. This implicated the role of RNA compaction in the formation of gel-like states, which translated to the appearance of irregular morphologies. Morphology changes were monitored at increasing temperatures, allowing us to visualize a possible mechanism by which rRNA relaxes at higher temperatures, resulting in liquid-like round shapes. Although temperature is a well-regulated parameter in many higher organisms, the variation in eukaryotic body temperatures between different species, and imposed by the environment, could give rise to different nucleoli dynamics. Our results provide an *in vitro* first step towards understanding the importance of temperature on the dynamics of RNA-based condensates.

An important caveat to consider is that our *in vitro* model is an amalgamation of components from different domains of life: the NPM1 protein is human, but the rRNA and 70S ribosomes are from *E. coli*. Although rRNA and ribosomes are relatively well conserved between the different branches of life, these molecules evolved to be stable in different Mg^2+^ and ATP concentration landscapes, and this can inevitably shape how they interact with one another. Nonetheless, this NPM1-rRNA model provides us with a good approximation of how these molecules will interact with one another as we expect RNA-RNA interactions to be relatively non-specific.

Ribosome assembly in the nucleoli is hypothesized to follow a gradient by which rRNA transcripts are made at the core of the nucleoli, followed by diffusion of rRNA through the organelle until the granular component, where it encounters the pre-ribosomal subunits. NPM1 proteins are localized to the granular component and facilitate the assembly of rRNA around the pre-ribosomal subunits. Once pre-ribosomal subunits are folded with their rRNAs, they are thermodynamically ejected from the condensate (11) before being shuttled to the cytoplasm where they undergo their final assembly. Pre-ribosomal subunits were recently observed to be clustered at the edges of the nucleoli using cryo-EM (41). Our results showing partitioning of the partially destabilized 70S ribosomes forming a halo around the rRNA-NPM1 condensates hint towards this spatial organization of ribosomes around the nucleoli. To our knowledge, this is the first time this has been shown *in vitro*. The 70S ribosome colocalization with NPM1 is also reminiscent of *in vivo* results where NPM1 was shown to interact with pre-ribosomal subunits (31). The decelerated fusion of the condensates after halo formation is reminiscent of recent work on how chromatin networks prevent the coalescence of nucleoli (42). The reversibility of this NPM1-ribosome interaction upon ATP depletion could be a feature of NPM1 function as some sort of multivalent shuttling protein that mediates ribosome transport through the nucleolus as well as the nucleus.

Together, our results not only help explain the discrepancies between *in vitro* and *in vivo* nucleoli dynamics, but also show that nucleoli are adaptive condensates, responding to local ATP concentrations through reversible Mg^2+^-induced compaction of the RNA component. Transient changes to ATP levels may be how cells regulate the amount of free Mg^2+^ and thereby control the dynamics of RNA-based condensates, such as the nucleolus.

## Materials and methods

For the complete experimental details, please refer to the Supplementary Information.

## Acknowledgments

This work was funded by the Netherlands Organization for Scientific Research (NWO-Startup to E.S.). We thank Richard Kriwacki for providing us with the NPM1-GFP plasmid, Davin Elian for his help in cloning the NPM1-wt plasmid. Lastly, we thank Aigars Piruska for his assiduous patience and microscopy technician skills.

## Supplementary Information

### Materials and Methods

#### Materials

Unless otherwise stated, all materials were obtained from Sigma-Aldrich.

#### Cloning of NPM1-wt into pET28a plasmid

The human NPM1 nucleic acid sequence was extracted by PCR from the pET28a(+)-NPM1-eGFP construct (gifted from R.K.) using the primers in **Table S1**, before being cloned into a pET28a(+) vector using restriction enzyme digest with *NdeI* and *XhoI*. The correct insertion was determined by sequencing (BaseClear, Leiden), and this construct called pET28a(+)-NPM1-wt.

**Table S1.**
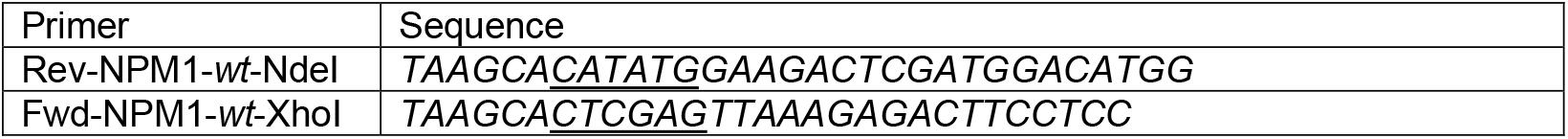

#### Protein expression, purification and labelling

*E. coli* BL21 (DE3) were transformed with pET28a(+)-NPM1-wt. Overnight cultures were used to inoculate large flasks of LB, then cells were grown at 37°C to an OD600 = 0.5-0.7, before protein expression was induced with 1 mM IPTG. Protein expression was carried out at 20 °C for at least 16 hours, after which the cells were harvested by centrifugation. The pellet was resuspended in lysis buffer (20 mM Tris-HCl, pH 7.5, 300 mM NaCl, 2 mM β-mercaptoethanol, 25 mM imidazole) containing 500 U Bezonase® Nuclease and Bovine Pancreas RNAse A (VWR). The resuspended cells were lysed using a homogenizer. Lysed samples were left at 4°C for 1 hour to allow the enzymes to degrade the nucleic acids. The lysate was spun at 35000 *xg*, 30 minutes at 4°C in a Beckman JA25.50 rotor. The clarified supernatant was loaded onto a 5 mL HisTrapFF (Cytiva). After loading, the column was washed with 50 mL of lysis buffer, and the His-tagged NPM1 proteins were eluted using elution buffer (20 mM Tris-HCl, pH 7.5, 300 mM NaCl, 2 mM β-mercaptoethanol, 250 mM imidazole). Eluted proteins were concentrated to <5 mL and loaded onto a Superdex 200 16/600 (GE Healthcare) size exclusion column connected to an AKTA Basic FPLC (GE Healthcare), and pre-equilibrated in storage buffer (20 mM Tris-HCl, pH 7.5, 300 mM NaCl). Fractionation of proteins was carried out at 1 mL/min, and monitored at 280 nm and 260 nm. The fractions of the main peak were pooled, and the protein concentration was determined using the NanoDrop One^C^ (Thermo Scientific). If the resulting UV spectrum had a 260/280 ratio of >0.6, the protein sample was further purified using anion exchange chromatography to remove contaminating nucleic acids. All protein isolates were dialyzed into storage buffer, aliquoted and snap frozen in liquid nitrogen and stored at -80 °C.

The NPM1 proteins were labelled using AlexaFluor488 C5 maleimide (Thermofisher Scientific) based on a previously published protocol (1). Excess dye was removed using anion exchange, and the protein dialyzed into storage buffer. The final protein concentration was determined using the NanoDrop One^C^. A labelled NPM1 stock was mixed with unlabeled protein at a 1:9 ratio of labelled:unlabelled protein, and this mixture was called NPM1-A488.

#### 70S ribosome purification

*E. coli* BL21 cells were harvested from 1L culture in LB media (A600 ∼1.5), and the cells were lysed using a bead beater in monosome gradient buffer (20 mM Tris-HCl, pH 8.0, 10 mM MgCl2, 140 mM KCl, 1 mM DTT, 0.1 mM EDTA, 20 U/mL s SUPERase•In™ RNase Inhibitor (Thermofisher). Cell debris was removed by centrifugation at 33000 *xg* for 30 minutes at 4 °C. The cleared supernatant was incubated at 37 °C for 80 minutes for ribosome run-off. The supernatant was again centrifuged at 33000 *xg* for 30 minutes at 4°C and additionally 0.22 μm filtered The ribosomes were pelleted by centrifugation for 3 hours at 48000 rpm at 4 °C in a Beckman Ti70.1 rotor. Pellets were gently dissolved in a small volume of monosome gradient buffer and the concentration was approximated using 1 A260 unit = 24 pmol of ribosomes per mL. Approximately 200 A260 units of crude *E*.*coli* BL21 ribosome isolate (∼0.5 mL) was adjusted to 1 mL with monosome gradient buffer and carefully loaded onto a 10-50% gradient of sucrose in monosome buffer in SW-28 tubes (for Beckman SW-28 rotor). The samples were centrifuged for 20 hours in a Beckman SW-28 rotor at 18000 rpm, 4°C. The resulting gradients were harvested at 4°C using a peristaltic pump and fractions of ∼1 mL were collected. Fractions containing the 70S ribosomes were pooled and pelleted by ultracentrifugation in the Ti70.1 rotor (Beckman) for 3 h at 48000 rpm, 4°C. The pellet was gently dissolved in 500 μL of monosome buffer, aliquoted in portions of 50 μL, flash frozen in liquid nitrogen and stored at -80°C.

#### 70S ribosome labelling

Sucrose gradient purified *E. coli* BL21 70S ribosomes were labelled as previously reported (2) using either ATTO488 NHS-ester (ATTO-TEC GmbH), or Dylight650 NHS-ester (Thermofischer Scientific) dyes. Briefly, 4.8 µM 70S ribosomes and 250 µM dye was mixed together in the reaction buffer (50 mM TrisHCl, pH 7.6, 15 mM MgCl2, 100 mM NH4Cl and 6 mM β-mercaptoethanol) and incubated at 37 °C for 30 minutes. Any precipitate was removed by centrifugation at 10000 *xg* for 1 minute using a tabletop centrifuge. The supernatant containing ribosomes was concentrated in a spin column (Vivaspin 6, MWCO 30 kDa) and thoroughly washed with reaction buffer to remove excess dye.

The final concentration of labelled 70S ribosome (A488) was estimated at 5.1 µM, and the ATTO488 label concentration to be 27.5 µM using a Nanodrop 1000 (Isogen), corresponding to ∼5-6 labels per ribosome. The labelled 70S ribosome (D650) was estimated at 6.3 µM, and the Dylight650 label concentration at 28.7 µM using a Nanodrop 1000 (Isogen), corresponding to ∼4-5 labels per ribosome. The ribosomes were then aliquoted, flash frozen in liquid nitrogen and stored at -80°C until further use.

#### *E. coli* rRNA purification

*E. coli* BL21 cells were harvested from a 1 L culture in LB media (A600 ∼1.5), and the cells were washed twice in buffer A (50 mM Tris, pH 7.7, 60 mM potassium glutamate, 14 mM magnesium glutamate, 2 mM DTT). Washed cells were lysed using the homogenizer, and insoluble debris was spun down at 20000 *xg* for 25 minutes. The supernatant containing the ribosomes was removed and the ribosomes were pelleted by centrifugation for 3 hours at 50000 rpm at 4°C in a Beckman Ti70.1 rotor. The pellet containing the ribosomes was resuspended in buffer A, and the rRNA was purified using standard phenol-chloroform extraction protocols. The final rRNA concentration was determined using the Nanodrop One^C^, where 1 OD260 = 40 μg/mL RNA.

#### RNA labelling using periodate oxidation

The 3’-hydroxyl of RNA was labelled with AlexaFluor™647-hydrazide (Thermofisher Scientific) using periodate oxidation. The RNAs labelled using this procedure included the purified rRNA, polyadenylic acid potassium salt (P9403 Sigma) or polyuridylic acid potassium salt (P9528 Sigma). Here, 2.5 nmol RNA in 100 mM sodium acetate (pH 5.2) was mixed with 2.5 mM freshly prepared sodium periodate to a total volume of 100 μL. This was left to incubate on ice for 50 minutes before nucleic acids were purified using either isopropanol precipitation followed by 70% ethanol precipitation, or using the Amicon®-Ultra spin concentrators (Millipore) (as per method described (3)). Either way, the RNA was mixed with 100 μL of 100 mM sodium acetate (pH 5.2) containing 25 nmol AlexaFluor647-hydrazide dye, and the sample was incubated at 4 for at least 24 hours in the dark. The labelled RNA was purified from the excess RNA by thoroughly washing it with several cycles of isopropanol and ethanol precipitation, or with several rounds of concentrating the sample and diluting it in Milli-Q water using Amicon®-Ultra spin concentrators. An agarose gel was used to double check free dye removal, before the sample concentrations were calculated using the Nanodrop One^C^, where 1 OD_260_ = 40 μg/mL RNA, and the dye fluorescence concentration was calculated using the extinction coefficient (at 649 nm) of 250000 cm^-1^M^-1^ (as described by the manufacturer). The AlexaFluor647-labelled RNAs were denoted with A647.

#### Making NPM1-RNA condensates

All experiments were performed using a standard base buffer (20 mM Tris-HCl, pH 7.2, 250 mM potassium glutamate) with different concentrations of magnesium glutamate (often denoted as Mg^2+^), as specified per experiment. The buffers were made as a 4X concentrated stock and were diluted to a final 1X working concentration, along with the other components, using Milli-Q water. The final concentrations of NPM1 protein and RNA used in each experiment were 20 μM and 100 ng/μL, respectively. In a typical experiment, the 4X buffer is first mixed with Milli-Q water, followed by RNA, then the NPM1, after which samples were mixed and pipetted onto a functionalized microscopy slide for imaging. Unless otherwise stated, samples were left to incubate on the glass slides at room temperature for at least 45 minutes before imaging.

#### Microscopy slides

Two types of microscopy slides were used: the 18-well Ibidi chambers for quick imaging, as well as PDMS chambers made in-house. The PDMS chambers were attached onto plasma-primed cover glass slides (No. 1.5H). These PDMS chambers were used for experiments that exceeded 1 hour, as Vaseline-sealed coverslips were applied on top to avoid evaporation. For both setups, the glass surfaces were cleaned using a plasma cleaner, then incubated for 1 hour with 0.1 mg/mL PLL-*g*-PEG dissolved 10 mM HEPES, pH 8.0, before the surface was thoroughly washed with Milli-Q water and dried with nitrogen gas.

#### Fluorescence confocal microscopy setups

Two fluorescence confocal microscopy setups were used for the experiments in this paper:

1. Olympus IX81 spinning disk confocal microscope, equipped with an Andor FRAPPA photobleach module and Yokogawa CSU-X1 spinning disk. The Andor 400 series solid state lasers were used to bleach and image the samples. All the images were recorded with a 100× oil immersion objective (NA 1.5) and an Andor iXon3 EM CCD camera.
2. Leica SP8 Liachroic-beam splitting confocal laser scanning microscope, equipped with a PMT detector, 2 x HyD SP GaAsP detectors and Leica DRC7000 GT monochrome camera. All the images recorded using this confocal used a HC PL APO 63x/1.40 (oil) CS2 (0.14 mm) objective.

#### Fluorescence recovery after photobleaching (FRAP)

Fluorescence recovery after photobleaching (FRAP) experiments were conducted on the Olympus IX81 spinning disk confocal microscope set-up. FRAP measurements were carried out by selecting a small ROI in the middle of a coacervate or aggregate of interest, and bleaching with the appropriate wavelength of the laser, depending on the sample. The 488 nm laser line was set at 100% laser power using 75 pulses of 150 μs, and the 647 nm laser line was set to at 100% laser power using 75 pulses of 100 μs. When both wavelengths were used, the 647 nm laser bleaching preceded that of 488 nm laser. The recovery was imaged at reduced laser intensity (at least 5–fold lower) and a regular time intervals, depending on the sample.

Using a MATLAB script, the experimental recovery was first normalized before being fitted to a simple exponential, as a first-order approximation of 2D diffusion with a fixed boundary (i.e. droplet edge) (4). Briefly, the exponential decay equation:

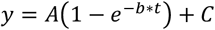

From this equation, the recovery half-life (τ) was calculated by *τ* = *ln*(2)/*b* and percentage recoveries were extracted by multiplying *A* by 100. We note that the theoretical maximum recovery is limited by the size of the bleached spot, which was 13.1 ± 4.4% of the droplet area in our experiments, corresponding to a maximum theoretical recovery of ∼87%.

#### Aging experiments

The CSU temperature stage (Tokai-HIT) was used to set the temperature of the samples. The sample temperatures achievable range from 8 – 39 °C, and is dependent also on the ambient temperature. Therefore, sample temperatures were always confirmed at the end of each experiment using a thermometer.

The condensate components and buffers were pre-incubated at the temperature of interest prior to being prepared using base buffer as described above. Samples were quickly mixed and immediately added to functionalized PMDS slides that were sealed with Vaseline. The FRAP experiments were performed over time, and subsequent analysis was performed (as described above) to determine τ.

An Arrhenius plot of the data revealed two straight lines with a transition around 21-22 °C (**Figure S2A**) and a transistivity plot (**Figure S2B**) was derived from the Arrhenius plot to show this drastic transition.

#### Circularity (Figures 2C and D)

The NPM1-rRNA condensates were mixed in either 5 mM Mg^2+^ or 10 mM Mg^2+^ buffer (as described above) and deposited onto the Vaseline-sealed PDMS slides incubating at 8 °C (sample temperature). Note the CSU temperature (Tokai-HIT) stage was set to 4 °C, but sample temperature was recorded throughout the experiment using a buffer blank in an adjacent well, and it is the sample temperature that is reported in our figures. Samples were incubated for at least 45 minutes before imaging began using the SP8 Leica confocal microscope. Sample temperature was adjusted incrementally with a 10-minute pause before imaging. At least three different images were captured and the circularities calculated here were averaged across the population of condensates.

To monitor changes to the condensate circularity, the NPM1-rRNA condensates were made in 14 mM Mg^2+^ buffer and left to incubate in Ibidi slides for 50 minutes before 11 mM ATP was added to a corner of the slide. The sample was imaged every 1 second using the Olympus IX81 confocal microscope, with a focus on one large condensate. The circularities calculated here were for this condensate over time.

Raw fluorescence confocal microscopy images and videos were processed and analyzed with MATLAB 2021 Image Processing Toolbox. Objects smaller than 200 pixels were excluded from the analysis (to avoid detecting smaller round droplets that tend to initially form as they settle onto the glass) and the area and mean intensity were extracted for every object in each frame. For circularity calculations, the perimeter of each object was calculated by taking the sum of all pixels that were directly adjacent to the dilute phase plus half the sum of all pixels that were only diagonally adjacent to the dilute phase. The circularity (*ϑ*) of the object cross section was then calculated as *ϑ* = 4 *πA/P*^2^, where *A* is the area of the object and *P* the perimeter (5). Subsequently, a single *ϑ* for each frame was calculated by taking the mean *ϑ* ± standard deviation.

#### ATP, ADP and AMP experiments (Figure 2 E-G)

Samples were prepared in a sequence as outlined above, and nucleotides were mixed last before everything was deposited onto a glass slide for imaging. The samples were mixed in base buffer containing 14 mM Mg^2+^, and each nucleotide was at 10 mM final concentration. The FRAP experiments were performed as outlined above. Errors are standard deviations from at least duplicate experiments.

#### SYBR Gold intensity calculations

SYBR Gold was used to indirectly probe rRNA compaction within the condensates. According to Kolbeck *et al*, the SYBR Gold 10000X stock is 12.4 ± 1.2 mM. Therefore, at 40 nM, the final SYBR Gold concentration used per 30 μL experiment is 62.5X below that of the reported 2.5 μM concentration where quenching effects were observed. SYBR Gold fluorescence is dependent on not just concentration, but also local environment (6). To test this, the condensate samples – using unlabeled components – were pre-mixed as outlined above and the SYBR Gold was added last. In some cases, the Mg^2+^ and ATP were added sequentially after condensate formation. There were no noticeable differences in fluorescence when comparing with pre-mixed and sequential samples. In fact, our reported results are a combination of sequential and pre-mixed intensities.

Since we were comparing SYBR Gold fluorescence between samples with different Mg^2+^ and/or ATP concentrations, the confocal images were taken using the Leica SP8 set-up. For all the images, the same laser settings were used. The fluorescence intensity of SYBR Gold from confocal microscopy images and videos were processed and analyzed with MATLAB 2021 Image Processing Toolbox. The images were binarized with an automatic intensity threshold computed with Otsu’s method (7). Objects smaller than 50 pixels were excluded from the analysis and the area and mean intensity were extracted for every object in each frame.

#### Partitioning experiments (Figure 3 A-C)

For partitioning of client molecules we prepared stock solutions at 33 μM 5,6-FAM-RP3 (CASLO) and 50 μM 5,6-FAM-SNORD52 (IDT) in the standard base buffer containing 5 mM Mg^2+^. The ribosome clients, 70S ribosomes-A488 and 70S ribosomes-D650, were thawed and centrifuged for 10 minutes at 20000 *xg* to remove any aggregates, and the supernatant was used for partitioning experiments. The ribosome concentration in the supernatants did not diverge much from the stock concentrations described above.

For each partitioning experiment, 1 μL of client was mixed with either the forming coacervates, or with the RNA component prior to NPM1 addition. Both orders of addition yielded similar results. The mixtures were incubated for 45 minutes on the glass slides at room temperature, before imaging.

For imaging, we used the Leica confocal setup with laser excitations of 488 nm (FAM, ATTO488) and 647 nm (Dylight650). For each laser setting, a blank was imaged made from NPM1-rRNA with one of the components labelled, accordingly.

The partitioning coefficient was then calculated using: *K*_*p*_ = (I_coacervate_ - I_background_)/(I_dilute_ - Ibackground), where Icoacervate is the average intensity as determined by a pixel gray value cut off above 40, and Idilute is the average intensity as determined by a pixel gray value cut-off below 10. The errors are standard deviations of at least three sets of *Kp* values derived from three images.

#### Ribosome halo formation

The ribosome halo formation was monitored over time using both confocal set-ups as highlighted in the video captions. The pixel intensity measurement (**Figure 3F**) was from a time-lapse experiment of ATTO488-labelled 70S ribosomes where the video was taken shortly after sample mixing. Using ImageJ, a line of fixed pixel length (equivalent to 9.23 μm) was drawn and the pixel intensity for each frame of the droplet was measured.

#### Apyrase enzyme experiments (Figure 6)

Apyrase Grade VII from Potato enzyme was dissolved in standard base buffer containing 5 mM Mg^2+^ (20 mM Tris-HCl, pH 7.2, 250 mM potassium glutamate, 5 mM magnesium glutamate). For the FRAP experiment, the samples were made using 5 mM Mg^2+^ base buffer with additives: ATP, AMP and ATP with apyrase. 5 U apyrase solution was added to the forming coacervate mix and all the samples were left to equilibrate on the Ibidi microscope slides for 30 minutes before imaging and FRAP experiments were performed. Error bars are standard deviations from at least two different photobleaching experiments.

The morphology changes to the NPM1-rRNA condensates were monitored using the Leica SP8 confocal microscope. Again, the samples were prepared as above, with 5U apyrase added to the forming coacervates. The sample was left to incubate on functionalized Ibidi slides for 30 minutes prior to imaging using both the 488 nm (NPM1-A488) and 647 nm (rRNA-A647).

Lastly, for the ribosome halo and apyrase experiments, 5 U apyrase solution was added to the formed coacervates on one side of the well of the microscope slide, and the disappearance of the ribosome halo was monitored every 1 second using the Olympus IX81 spinning disk confocal microscope set-up.

## Supplementary Figures

**Figure S1.**
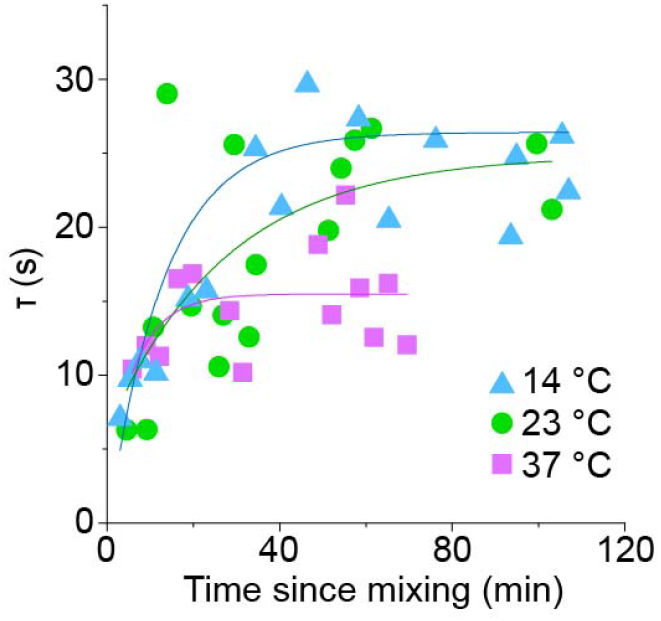
The τ of NPM1 remains relatively constant over time for aging NPM1-rRNA droplets in 0 mM Mg^2+^ buffer at 14°C, 23°C and 37°C. The lines are exponential fits of the individual measurements, from which the asymptote value and its error were plotted in Figure 2B in green.

**Figure S2.**
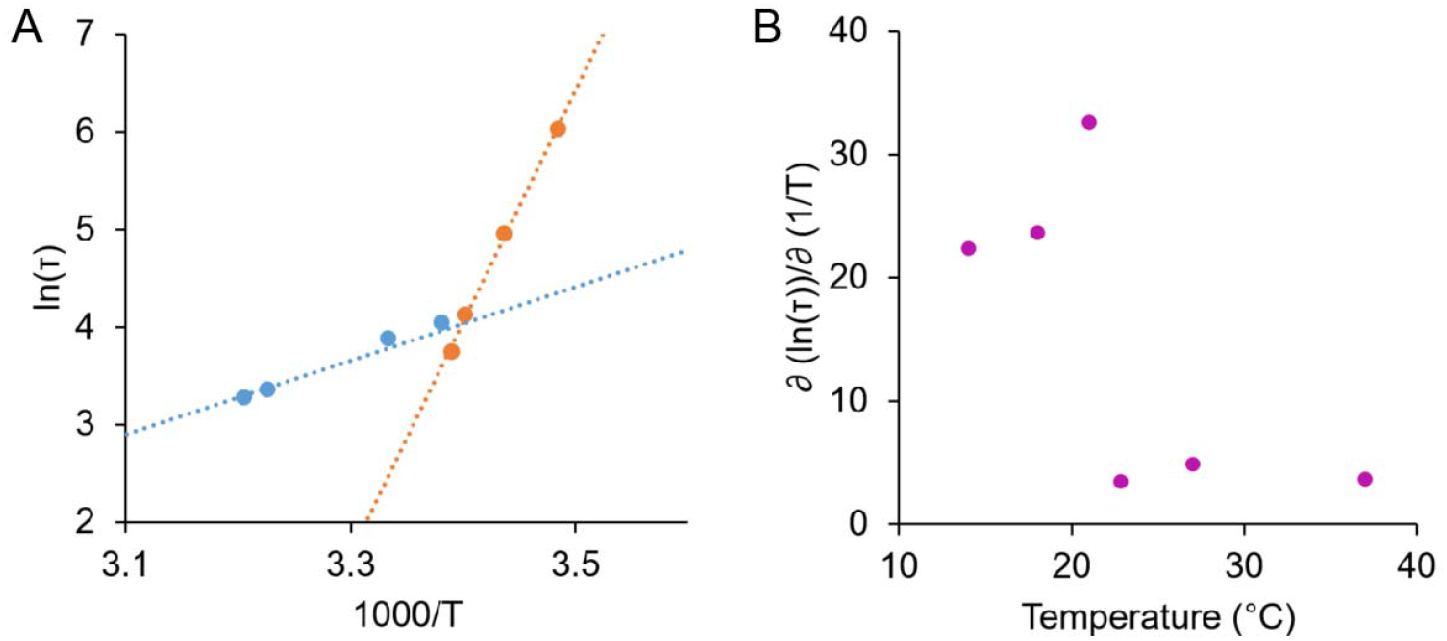
Arrhenius equations (**A**) and transistivity plot (**B**) for aging rRNA component of NPM1-rRNA condensates made in 0 mM Mg^2+^ buffer. This shows the Arrhenius equation is constant above 22°C, and increases drastically below 22°C.

**Figure S3.**
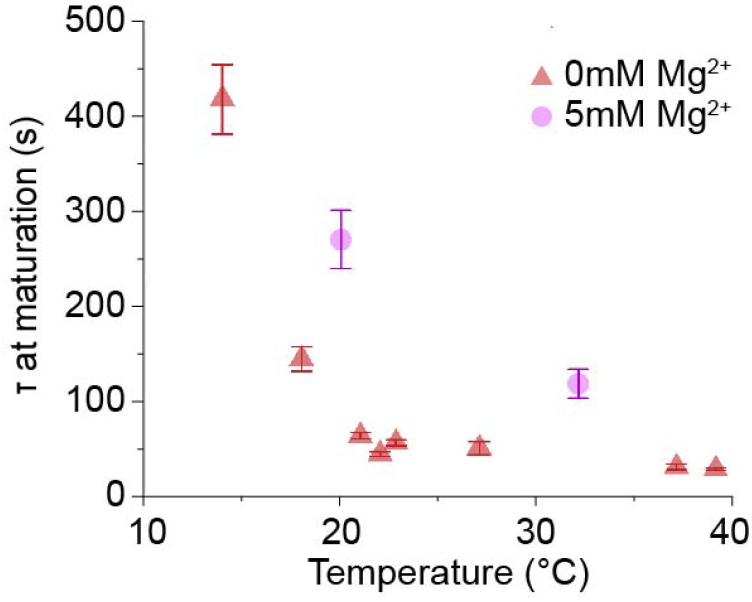
Slower τ for the rRNA component was observed at 5 mM Mg^2+^ than at 0 mM Mg^2+^. The errors were derived from the exponential fits to the τ data monitored over time.

**Figure S4.**
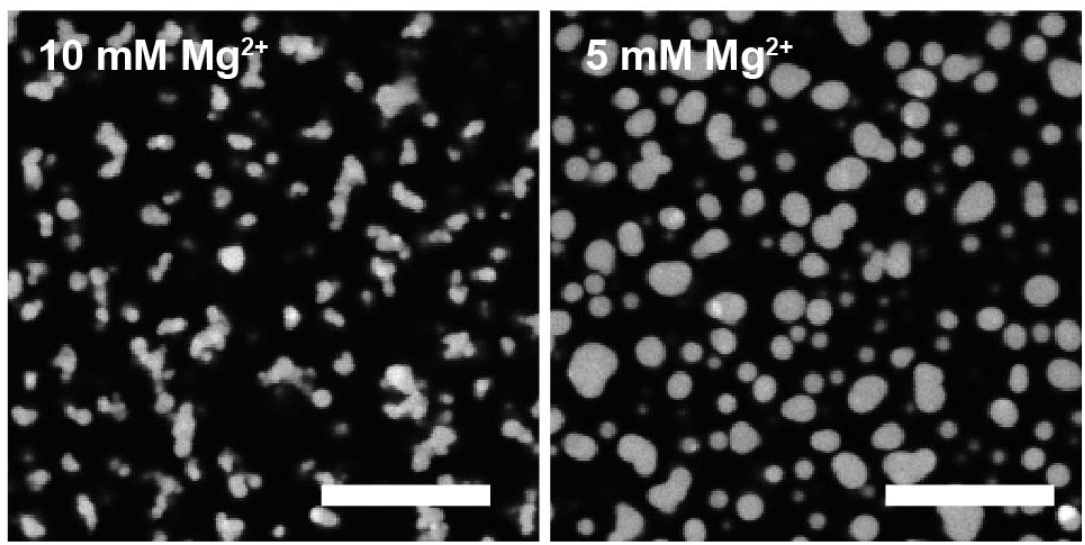
At 8°C, NPM1-rRNA condensates made at 10 mM Mg^2+^ appeared smaller and had lower average circularity than condensates made in 5 mM Mg^2+^ buffer (see Figure 2C). Scale bars are 10 μm.

**Figure S5.**
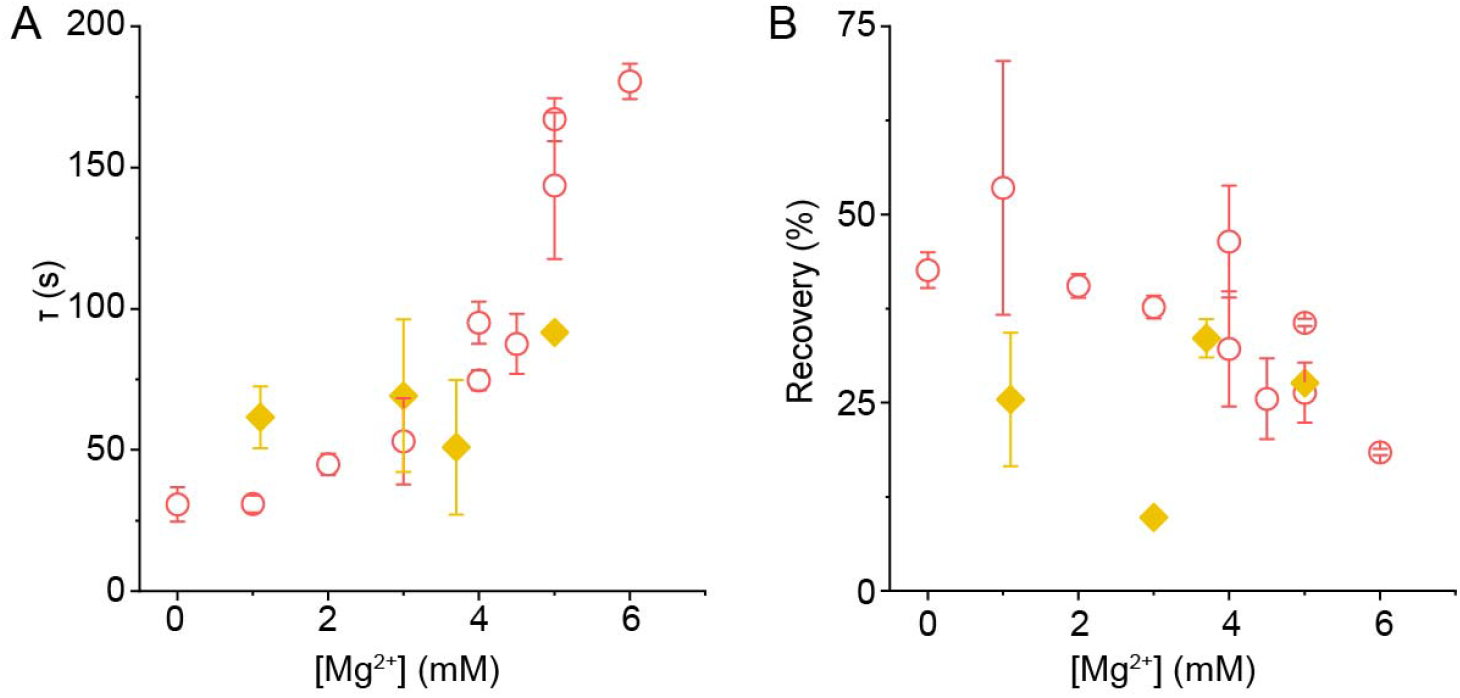
The FRAP parameters for the rRNA component, τ (**A**) and percentage recovery (**B**), in samples after ATP addition (yellow) at different final Mg^2+^ concentrations. These results are overlaid with the parameters found when only increasing Mg^2+^ in the buffer (red – as in Figure 1D). The errors represent standard deviations from at least three replicates.

**Figure S6.**
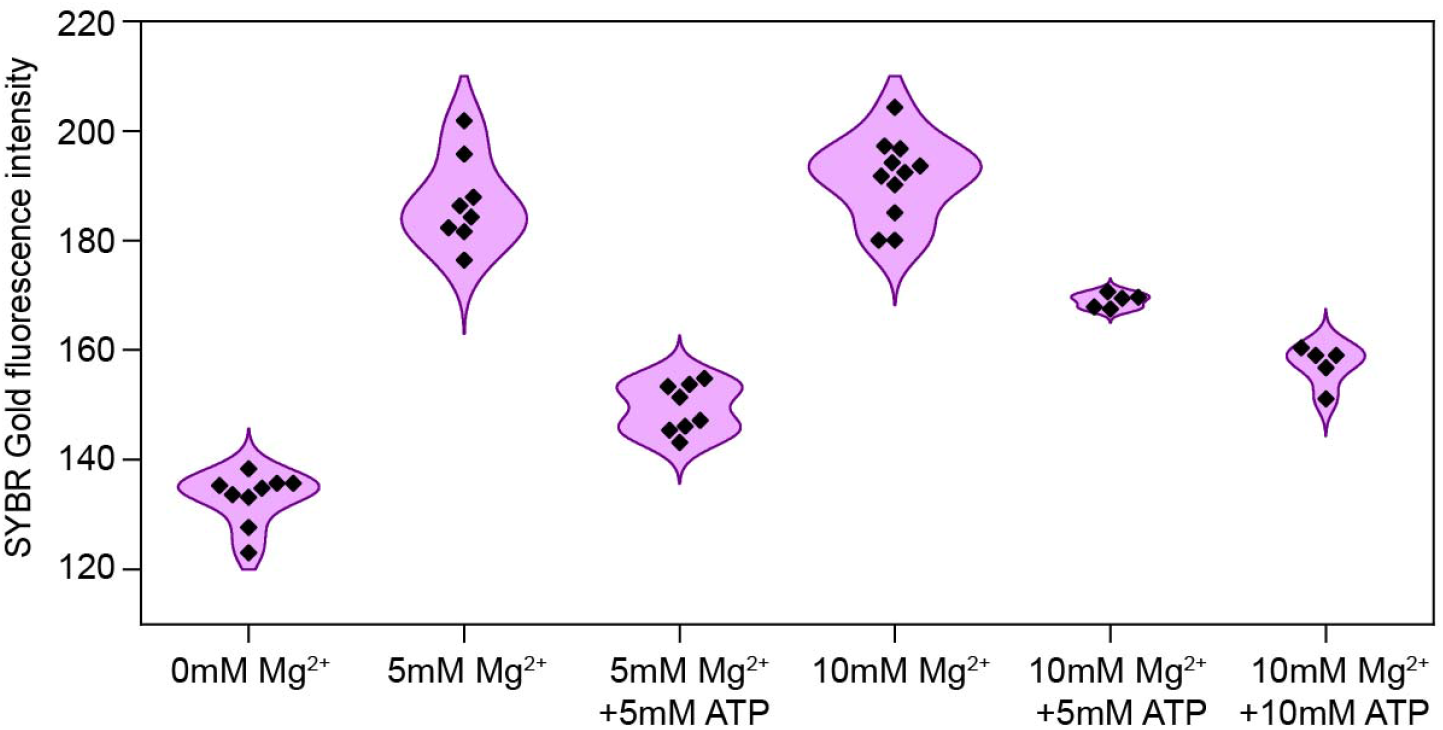
SYBR Gold fluorescence intensity for a population of condensates. Each point is the averaged intensity from a single confocal image, which shows the distribution of fluorescence intensities inside the NPM1-rRNA condensates. This result demonstrates that Mg^2+^ increases the SYBR Gold fluorescence intensity, which would correlate to increased RNA compaction, whereas ATP addition reduces SYBR Gold fluorescence, and also RNA compaction.

**Figure S7.**
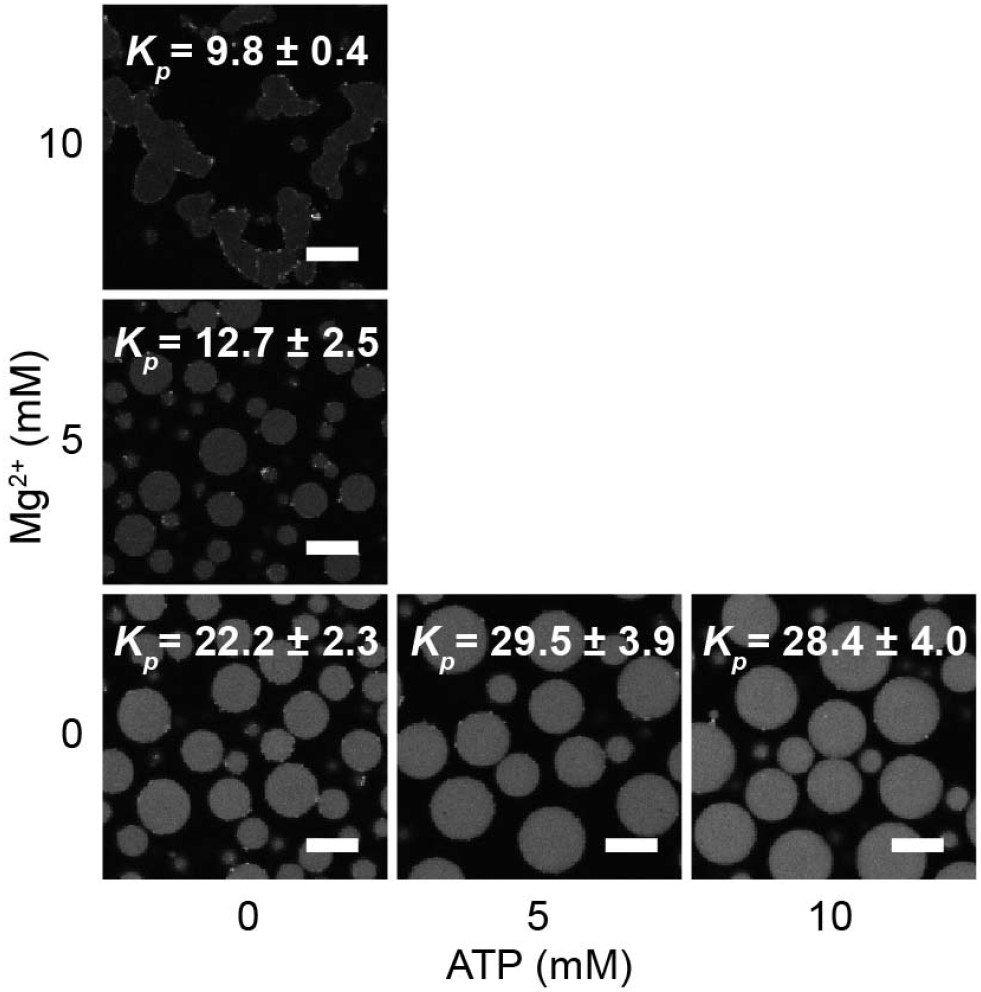
Ribosome partitioning at different Mg^2+^ concentrations and ATP concentrations inside NPM1-rRNA condensates. The partitioning coefficient (*Kp*) errors were derived from an average of values from at least four different images. Scale bars are 10 μm.

**Figure S8.**
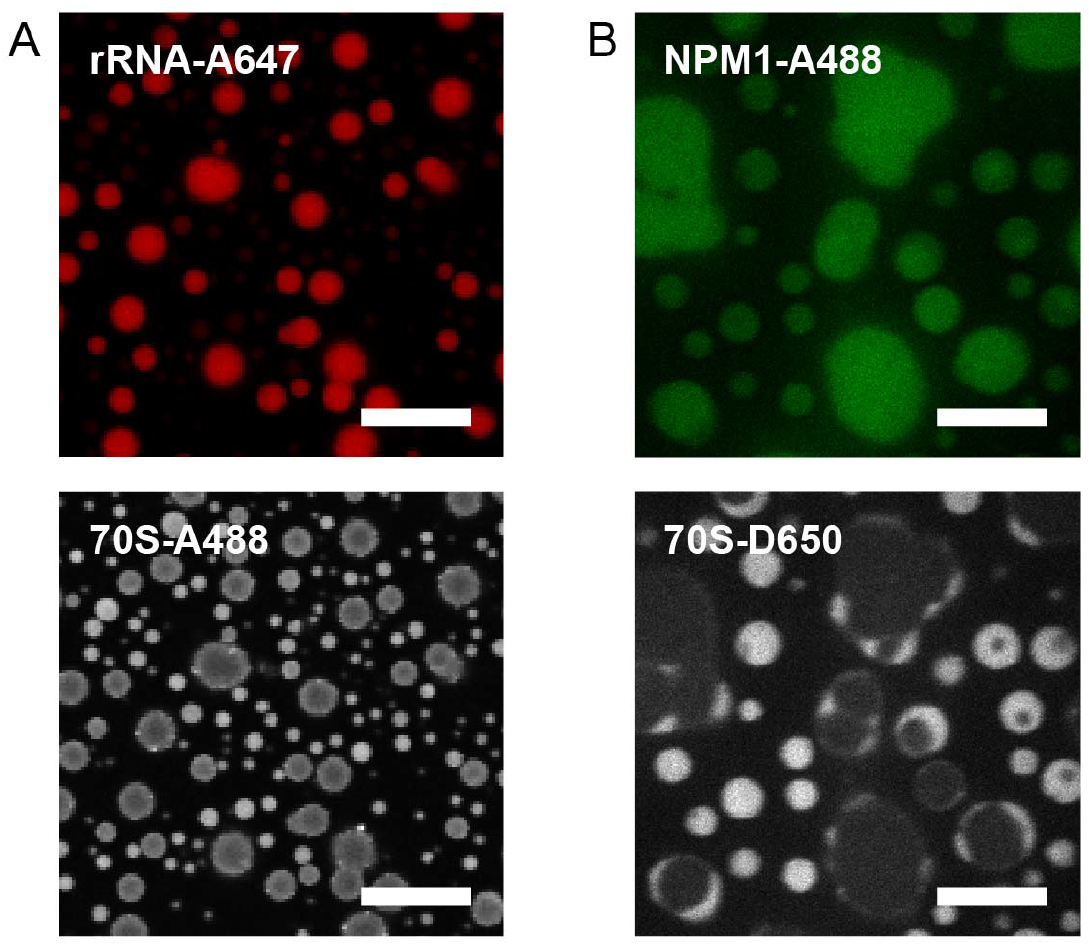
Confocal images showing the formation of the ribosome halo upon 5 mM ATP addition to pre-formed NPM1-rRNA condensates in 10 mM Mg^2+^ buffer. (**A**) Images from two different wavelengths of the same frame, showing 70S ribosomes forming a halo around the rRNA-647 condensates, as well as new droplets where rRNA is excluded from the ribosome fluorescence. These images are the last frame of Video S4. (**B**) Images from two different wavelengths of the same frame, showing NPM1 fluorescence co-localizing with 70S ribosomes in the halo as well as in droplets. Scale bars are 10 μm.

## Supplementary Video Captions

**Video S1**. NPM1-rRNA condensates formed in 5 mM Mg^2+^ buffer showing slowed droplet fusion in the order of minutes. This video is a series of images taken using the SP8 Leica confocal microscope with the 488 nm laser showing the NPM1-A488 fluorescence.

**Video S2**. NPM1-pU condensates at 20 mM Mg^2+^ shows round morphology and <1s fusion. This video is a series of images taken using the Leica confocal microscope with the 647 nm laser showing the pU-A647 fluorescence.

**Video S3**. Ribosome halo formation as NPM1-rRNA coacervates coalesce, with decelerating fusion and puncta observed. This video is a series of confocal images taken using the Olympus IX81 with the 488 nm laser illuminating the ATTO488-labelled 70S ribosomes.

**Video S4 A & B**. NPM1-rRNA condensates that were pre-formed at 10 mM Mg^2+^ prior to 5 mM ATP addition. These two videos are of the same sample after ATP addition at two different wavelengths (**A** 488 nm laser for 70S ribosome-A488, and **B** 647 nm laser for rRNA-A647). The ribosome halo forms around the edge of the NPM1-rRNA condensates, with also external ribosome-NPM1 droplets appearing after 10 minutes.

**Video S5**. Upon apyrase addition, the ribosome halo around NPM1-rRNA condensates dispersed. This video is a series of confocal images taken using the Olympus IX81 with the 647 nm laser illuminating the Dylight650-labelled 70S ribosomes.

**Video S6**. Upon apyrase addition, the ribosome halo around NPM1-rRNA condensates as well as internal puncta disappeared. The disappearance of the ribosome halo also results in the relaxation of the NPM1-rRNA causing them to fuse together. This video is a series of confocal images taken using the Olympus IX81 with the 647 nm laser illuminating the Dylight650-labelled 70S ribosomes.

